# Genome wide association with quantitative resistance phenotypes in *Mycobacterium tuberculosis* reveals novel resistance genes and regulatory regions

**DOI:** 10.1101/429159

**Authors:** Maha R Farhat, Luca Freschi, Roger Calderon, Thomas Ioerger, Matthew Snyder, Conor J Meehan, Bouke de Jong, Leen Rigouts, ALex Sloutsky, Devinder Kaur, Shamil Sunyaev, Dick van Soolingen, Jay Shendure, Jim Sacchettini, Megan Murray

## Abstract

Drug resistance is threatening attempts at tuberculosis epidemic control. Molecular diagnostics for drug resistance that rely on the detection of resistance-related mutations could expedite patient care and accelerate progress in TB eradication. We performed minimum inhibitory concentration testing for 12 anti-TB drugs together with Illumina whole genome sequencing on 1452 clinical *Mycobacterium tuberculosis* (MTB) isolates. We then used a linear mixed model to evaluate genome wide associations between mutations in MTB genes or noncoding regions and drug resistance, followed by validation of our findings in an independent dataset of 792 patient isolates. Novel associations at 13 genomic loci were confirmed in the validation set, with 2 involving noncoding regions. We found promoter mutations to have smaller average effects on resistance levels than gene body mutations in genes where both can contribute to resistance. Enabled by a quantitative measure of resistance, we estimated the heritability of the resistance phenotype to 11 anti-TB drugs and identify a lower than expected contribution from known resistance genes. We also report the proportion of variation in resistance levels explained by the novel loci identified here. This study highlights the complexity of the genomic mechanisms associated with the MTB resistance phenotype, including the relatively large number of potentially causative or compensatory loci, and emphasizes the contribution of the noncoding portion of the genome.

## Introduction

Tuberculosis (TB) remains a major global public health threat. In 2016 there were an estimated 10.4 million TB cases globally and 1.7 million deaths due to the disease. One of the most challenging forms of disease is caused by multidrug resistant (MDR) *Mycobacterium tuberculosis*, with a global annual incidence of over half a million cases^1^. The World Health Organization (WHO) estimates that only two of every three patients with multidrug resistant TB are diagnosed, three in every four of the diagnosed are treated, and only one of every two of the treated patients are cured, resulting in the grim reality of about 75% of the incident cases persisting in the community or succumbing to their illness. Antibiotic resistance is also an increasing problem in other human pathogens, and transmission of antibiotic resistance from person to person is amplifying the public health threat^2^.

Improved surveillance, diagnosis and treatment are designated priorities by the WHO and the US, European CDCs for addressing the antibiotic resistance challenge^1,3,4^. These measures will rely on an improved understanding of the mechanisms of resistance acquisition in bacteria. The knowledge of genetic mechanisms of antibiotic resistance has formed the basis of several commercial molecular diagnostics for TB that have had remarkable global uptake, despite the fact that they only reliably test for a subset of TB drugs and hence have not yet been able to replace the traditional more costly and slow process of mycobacterial culture and drug susceptibility testing (DST)^1,5–7^. Understanding antibiotic resistance mechanisms and methods that compensate for lost bacterial fitness in the context of antibiotic resistance can also pave the way for the development of companion drugs that restore antibiotic susceptibility^8,9^ and can open the possibility of ‘evolutionarily directed’ therapies that can aid in primary prevention of resistance acquisition^10^.

To date, attempts at genome wide association for antibiotic resistance in *Mycobacterium tuberculosis* (MTB) have been limited by the relatively low number of isolates phenotypically resistant to antibiotics, and have exclusively relied on phenotypes defined by drug susceptibility testing (DST) performed at a single ‘critical concentration’, likely a result of convenience sampling from clinical isolate archives in clinical mycobacterial laboratories^11–13^. Although such ‘binary’ DST is currently the standard to guide patient care, MTB critical concentrations are largely based on consensus and lack solid scientific support. The WHO has also declared that “the critical concentration defining resistance is often very close to the minimum inhibitory concentration required to achieve anti-mycobacterial activity, increasing the probability of misclassification of susceptibility or resistance and leading to poor reproducibility of DST results”^14^. Although more laborious and expensive, the quantification of the resistance phenotype through minimum inhibitory concentration (MIC) testing is considered a major improvement in the current standard for clinical phenotyping of drug resistance^15^, and MICs are more appropriate for the assessment of the biological effects of genomic variation in understanding the mechanism of resistance and bacterial fitness. The association of this variation with MICs also promises to refine our molecular prediction of antibiotic resistance for clinical and diagnostic use, as considerable gaps remain in prediction of resistance to first line drugs like pyrazinamide (PZA), ethambutol (EMB) and second line drugs^16,17^. Here we present a study of 1526 isolates where MICs were measured for 12 anti-tubercular agents and whole genome sequencing and genome wide association was performed. We also validate our findings in a globally representative public set of TB genomes with binary DST phenotypic data.

## Results

Of the total 1526 isolates included in the primary analysis, 76 isolates were excluded because their sequencing data did not meet coverage and mapping criteria (methods). The remaining 1452 isolates originated from 24 different countries, but the majority, 1,226, was from Peru. The isolates were each tested against a minimum of four and up to 19 drugs with a median of 12 drugs/isolate (Table S1). Figure 1A provides histograms of the MIC results for isoniazid (INH), PZA, amikacin (AMI) and moxifloxacin (MXF) (complete set of histograms in Figure S1). Overall, 976 isolates were MDR (INH MIC >0.2mg/dl & rifampicin (RIF) MIC >1mg/dl) and 438 were pre-XDR (i.e. additionally resistant to either a fluoroquinolone, MXF, ciprofloxacin (CIP) or ofloxacin (OFX) or a second line injectable, SLI *i.e*. capreomycin (CAP), kanamycin (KAN) or AMI. A total of 157 isolates were XDR, *i.e*. MDR and resistant to a fluoroquinolone *and* a SLI. Despite testing at multiple concentrations close to the critical cutpoint in this sample enriched for MDR, we observed a low rate of intermediate MICs for most first and second line agents with notable exceptions for the drugs EMB, PZA, streptomycin (STR) and ethionamide (ETA) (Figure 1A & Figure S1).

**Figure 1 A:**
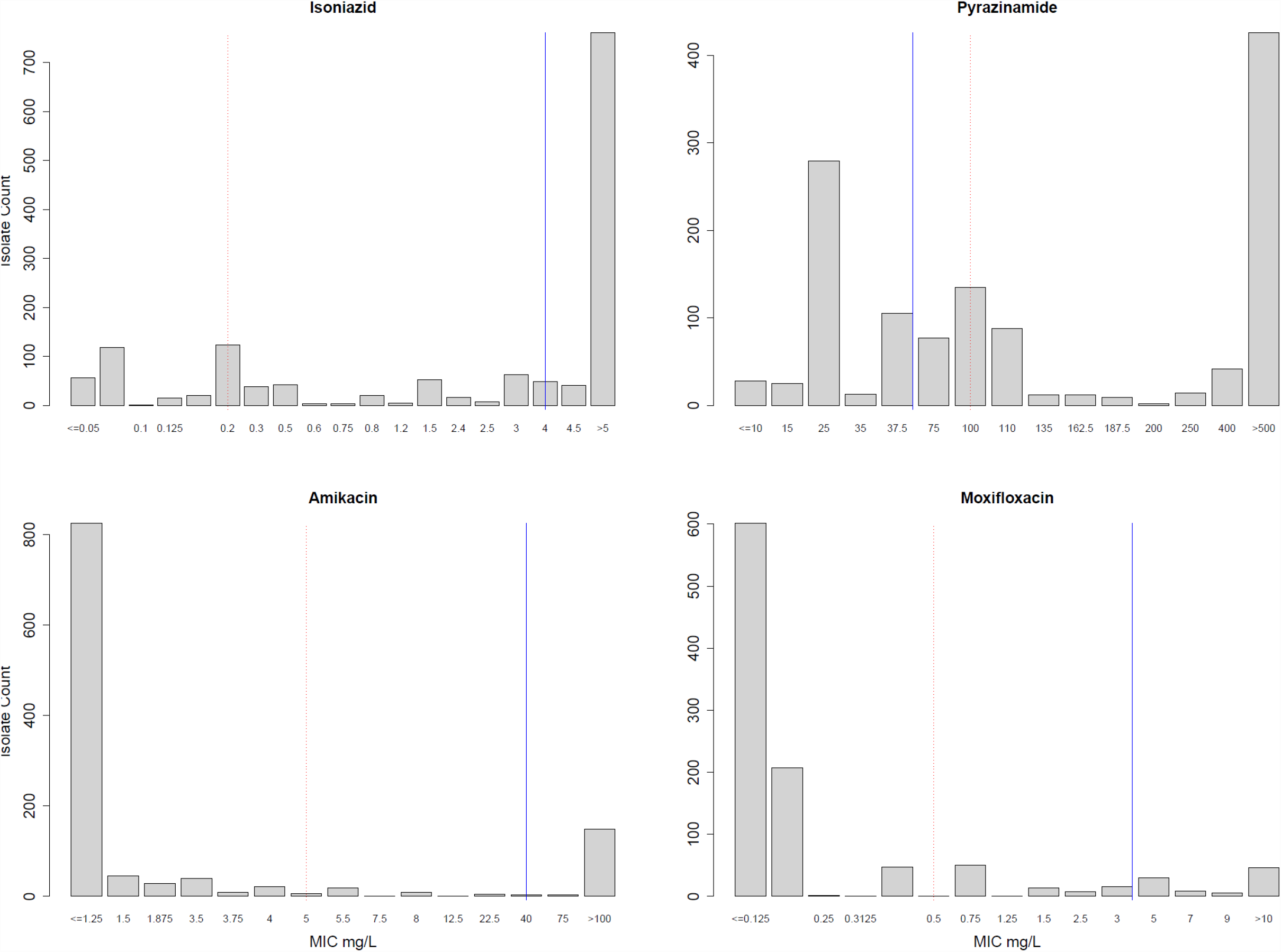
MIC distributions for 4 drugs. Dotted red line represents the WHO recommended critical concentration on 7H10 media, and the blue line represents the lower limit of achievable serum concentration from pharmacodynamic studies (Table S7).

We identified 73,778 unique genetic variants in the 1,452 genomes. The majority of the variants, 42,871 (58%) occurred in only one of the 1,452 isolates (Figure 1B) and the majority of single nucleotide substitutions (SNVs) in coding regions were nonsynonymous amounting to 36,479 vs 20,541 that were silent. We identified 7,178 variants with a frequency of >0.01 of which 2,701 had a frequency of >0.05. In addition to SNVs we observed an appreciable number of insertions and deletions (indels), with 9% of the observed variants with an AF >0.05 being indels. Furthermore, the noncoding portion of the genome (10.3% by length) harbored a slightly disproportionate degree of variation with 13% of SNVs with an AF>0.05 occurring in these regions.

**Figure 1 B:**
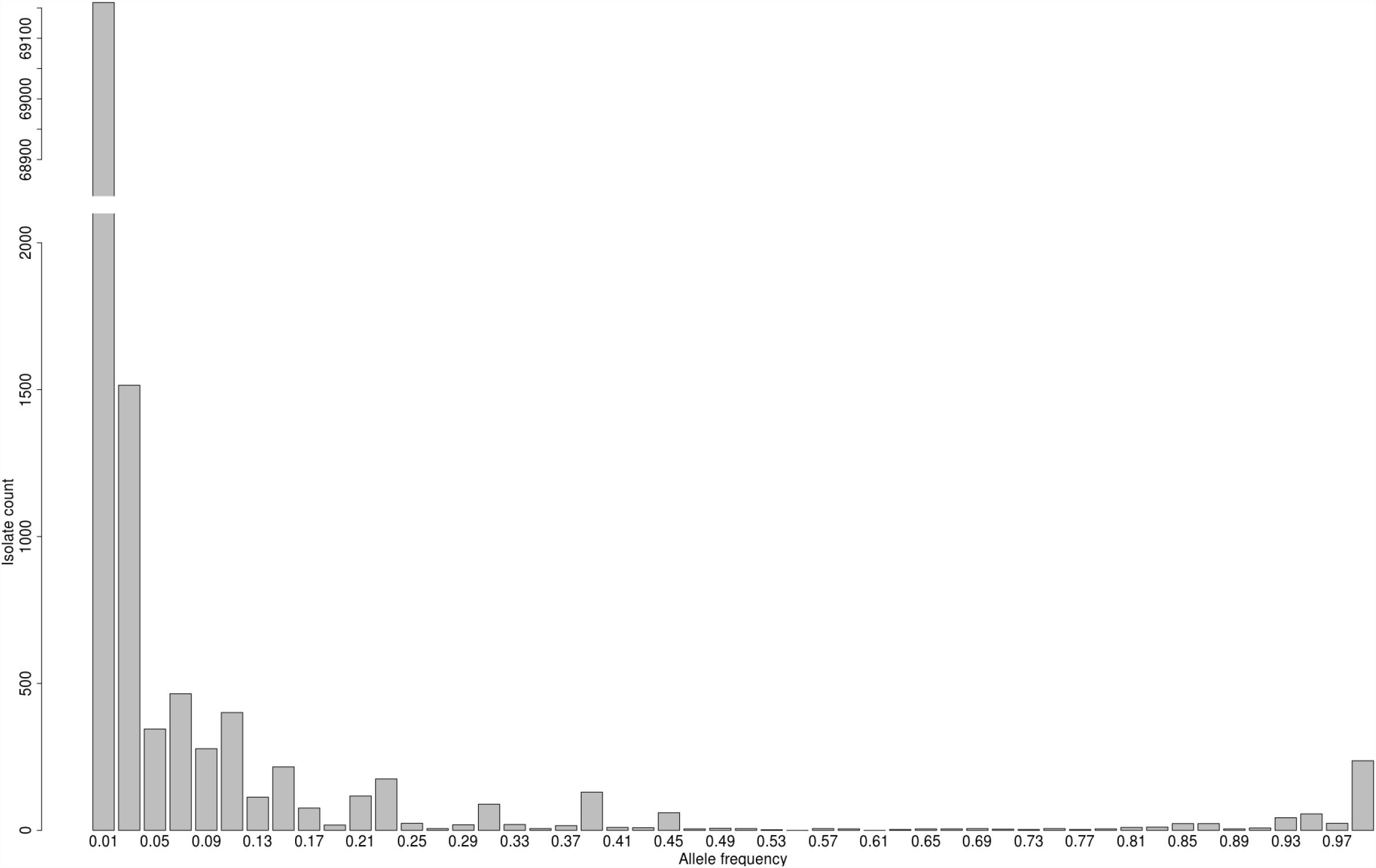
Allele Frequency distribution relative to H37Rv. Frequencies in the text given as minor allele frequencies, i.e. folded allele frequencies relative to H37Rv. The isolate count axis interrupted and scaled to accommodate the large

The isolates’ lineage diversity was consistent with their geographic origin with 86% being lineage 4 but diverse within this lineage with 39% of the total being lineage 4.3 (LAM), 31% lineage 4.1 (Haarlem) and 16% representing other L4-sublineages. Of the total 11% belonged to Lineage 2. There were a total of 43 isolates that belonged to other lineages (L1, L3 & L5). Figure 1C displays the pairwise genetic covariance between the isolates, and demonstrates that although the majority were lineage 4 there was considerable diversity among the isolates.

**Figure 1 C:**
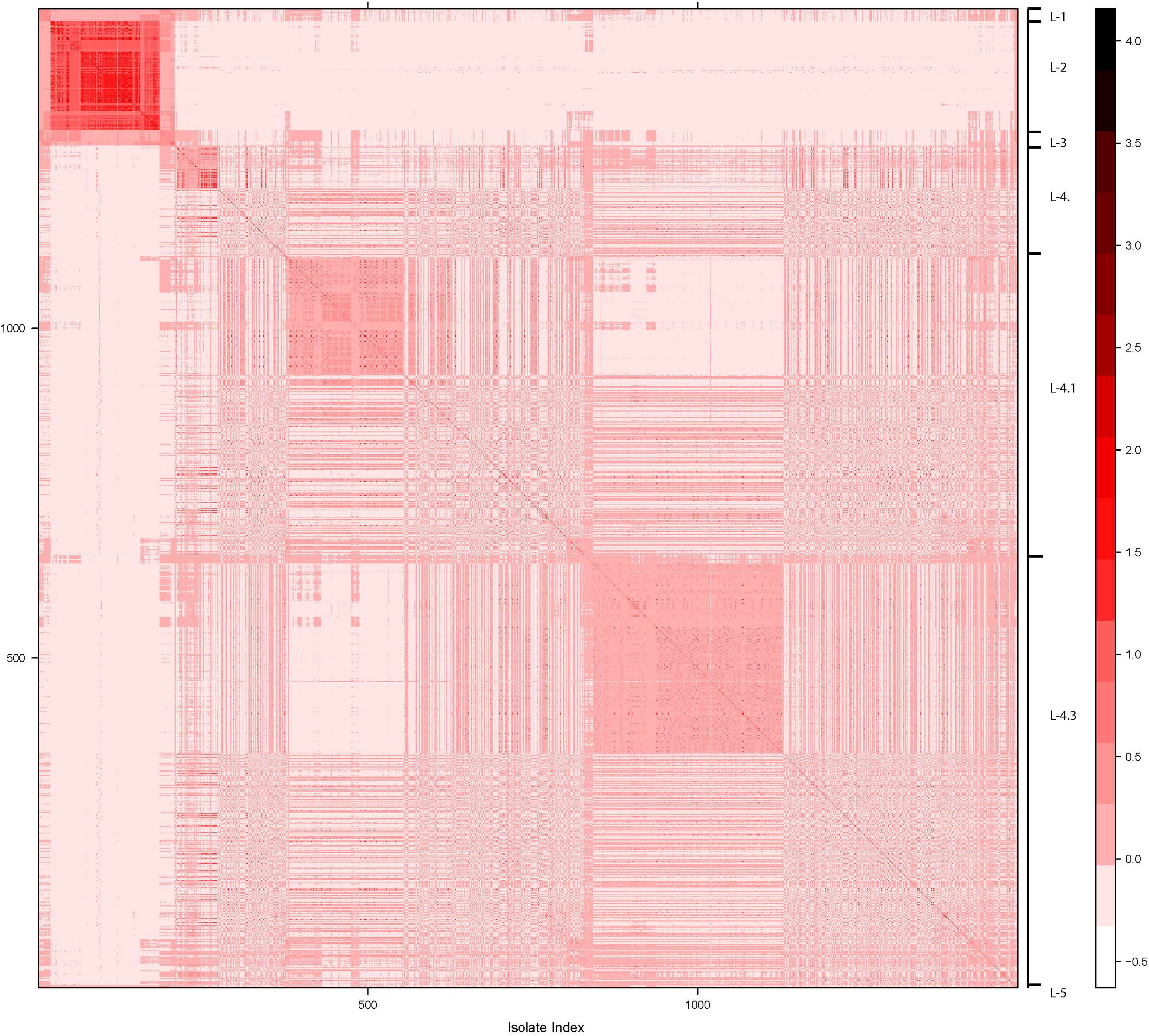
Heatmap displaying genome level similarity of the isolates used for the test GWAS. Similarity was measured using the isolate-isolate genetic covariance. Darker red indicates higher similarity/covariance. Attached separately due to size.

Genome wide association was performed for each drug separately using a gene/noncoding region binary burden score, excluding any loci with burden frequency of < 0.01, and correcting for population structure by fitting a linear mixed model. A total of 2791 loci had a burden frequency of ≥0.01. We set the significance threshold at an FDR <0.05 as we planned to perform validation on an independent dataset. QQ plots of the resultant p-value distribution suggested that the correction for population structure was adequate (Figure S2). Twenty known resistance loci (methods) were identified by genome-wide association and for all drugs known loci were associated with the highest effect size and lowest P-value of all the significant hits (Table S2). The RNA polymerase β-subunit gene *rpoB* was the most significant hit across all drugs with a RIF logMIC increase of 3.24 log(mg/L) and P-value of <10^-187^. Of the known locus-drug associations detected, the smallest effect size was measured for the *embA-embC* intergenic region, an EMB logMIC increase of 0.45 at a P-value of 1×10^-7^. Notably we did not identify a significant association between the compensatory gene *rpoA* and RIF resistance, the *embA*& *embC* genes and EMB resistance and between *gyrB* and MXF resistance. Given stepwise and co-linear development of antibiotic resistance in MTB and the prevalence of MDR in our sample, most of the known resistance loci were identified to be associated with more than one antibiotic, but in each case the known causative locus was the most significantly associated with its respective drug (Table 1, Table S2). We implicated several promoter/intergenic regions surrounding known genes including not only the *Rv1482c-fabG1* and the *eis-Rv2417c* intergenic regions that are currently used in one or more commercial diagnostics^6,25^, but also the regions upstream of *embAB (embA-embC), pncA (pncA-Rv2044c)*, and *ahpC (oxyR’–ahpC)*. The known compensatory gene *rpoC* was strongly associated with resistance to both RIF and rifabutin. We also identified the *rpsA* gene to be associated with PZA resistance with an effect size and P-value lower that of variants in the intergenic region containing the *pncA* promoter (0.55 logMIC increase & 2×10^-4^ vs 0.81 & 7×10^-5^ respectively, Table S2).

**Table 1:** Novel regions that were confirmed in the second validation GWAS. All drugs to which they were found to be associated are listed. The first drug listed was the drug found to be most significantly associated and for which the GWAS results are listed in the subsequent columns. For reference 6 known resistance loci are listed at the end along with their respective allele frequency, effect size and P-value. Full results are detailed in Tables S2–3. Drug abbreviations detailed in Table S7. OR: odds ratio. SE standard error. *transmembrane protein, **integral membrane transport protein, ***probable methyl transferase & membrane protein, ^transcriptional regulator, †structure specific helicase.

| Locus/site | Test GWAS (n=1452) |  |  |  |  | Validation GWAS (n=792) |  |  |  |  |
| --- | --- | --- | --- | --- | --- | --- | --- | --- | --- | --- |
|  | Drug | Allele frequency | Scaled Effect size (logMIC) | Scaled SE (logMIC) | P-value raw | Drug | Allele frequency | OR | SE | P-value raw |
| *ubiA* (Rv3806c) | EMB, INH, RIF, PZA, KAN | 0.071 | 0.52 | 0.07 | 1E-13 | EMB, INH, RIF, PZA | 0.066 | 1.25 | 0.10 | 3.6E-03 |
| *Rv3805c* - 4267647T>C (D397G) | AMI | 0.050 | 2.72 | 0.63 | 4E-06 | AMI | 0.283 | 1.11 | 0.03 | 4.7E-04 |
| *sirA* | ETA | 0.070 | 0.78 | 0.21 | 1E-04 | CAP, KAN, STR | 0.015 | 1.27 | 0.09 | 8.0E-04 |
| *whiB6*<sup>^</sup> | CAP, AMI, KAN | 0.037 | 0.59 | 0.15 | 3E-05 | CAP, AMI | 0.069 | 1.16 | 0.06 | 2.7E-03 |
| *ccsA* | KAN | 0.052 | 1.64 | 0.47 | 3E-04 | AMI, CAP | 0.274 | 1.11 | 0.03 | 2.7E-04 |
| *RNase J* (Rv2752c) | INH, RIF | 0.042 | 0.77 | 0.20 | 8E-05 | KAN, AMI, RIF, INH | 0.067 | 1.29 | 0.06 | 2.8E-08 |
| *PPE35* | PZA | 0.112 | 0.54 | 0.14 | 1E-04 | EMB | 0.334 | 1.24 | 0.08 | 4.9E-04 |
| *Rv3434c* * | KAN, AMI | 0.048 | 1.14 | 0.33 | 1E-04 | KAN, AMI | 0.047 | 1.18 | 0.06 | 2.3E-03 |
| *thyX-hsdS.1* | AMI | 0.018 | 0.74 | 0.22 | 3E-04 | STR | 0.016 | 1.32 | 0.13 | 3.9E-03 |
| *dinG*<sup>†</sup> | RIF | 0.069 | 1.86 | 0.47 | 2E-05 | AMI, STR | 0.293 | 1.10 | 0.03 | 8.8E-04 |
| *espK-espL* | RFB, RIF | 0.053 | 1.21 | 0.34 | 2E-04 | AMI, STR | 0.283 | 1.11 | 0.03 | 4.7E-04 |
| *kefB* (Rv3236c)** | PZA | 0.137 | 0.80 | 0.24 | 6E-04 | RIF, INH, PZA, EMB, STR | 0.235 | 1.60 | 0.16 | 2.0E-06 |
| *Rv2952* *** | EMB | 0.067 | 0.64 | 0.17 | 2E-04 | STR, AMI, INH, CAP | 0.203 | 1.29 | 0.08 | 2.6E-05 |
| *inhA* | INH | 0.03 | 1.10 | 0.26 | 2E-05 | - | - | - | - | - |
| *Rv1482c-fabG1* | INH | 0.10 | 0.63 | 0.16 | 6E-05 | - | - | - | - | - |
| *pncA* | PZA | 0.24 | 1.40 | 0.07 | 4E-89 | - | - | - | - | - |
| *pncA-Rv2044c* | PZA | 0.02 | 0.81 | 0.20 | 7E-5 | - | - | - | - | - |
| *embB* | EMB | 0.28 | 0.94 | 0.04 | 9E-101 | - | - | - | - | - |
| *embC-embA* | EMB | 0.03 | 0.45 | 0.09 | 1E-7 | - | - | - | - | - |

We identified 50 novel loci to be associated with resistance to one or more antibiotics (Table S2). Sixteen loci were associated with resistance to more than one drug. Two such loci were associated with resistance to all three SLI agents, the gene encoding the transcriptional regulator WhiB6, the cytochrome P450 oxidoreductase encoding *fprA* gene (logMIC change & P-value: 0.59 & 1×10^-4^, 1.37 & 1×10^-6^ respectively). CcsA a gene in the cytochrome P450 maturation pathway was also associated with SLI resistance (KAN logMIC change 1.64 & P-value 2×10^-4^ –Tables 1 & S5) with an effect size among the top 10 measured for the novel loci. The most significantly associated novel locus was the gene *ubiA* (Rv3806c) with the drug EMB (logMIC 0.52 & P-value 1×10^-13^). The locus *Rv3083* which encodes the gene *mymA*, an alternative monoxygenase to *ethA*^41^, was associated with resistance to ETA and two other drugs and was among the 10 most significant novel hits (ETA logMIC 0.60 & P-value 1×10^-4^). Twelve intergenic regions were found to be associated with resistance including the intergenic regions *thyX-hsdS.1* and *glnE-glnA2*, as well as regions adjacent to type VII secretion system related genes like *espK-espL* (Table 1 & Table S2). The secondary genome wide association performed at the site level identified associations of individual substitutions (SNV) or indels within the loci associated in the primary analysis (Table S3). In addition, four SNVs in other novel loci: L111M in *Rv3327*, D397G in gene *aftB*, 3778221GA in the intergenic region *spoU*-*PE-PGRS51*, and 640954AG in the intergenic regions *Rv0550c-fadD8* were associated with resistance (Table S3). No novel associations were found for the drug linezolid.

The 50 novel associations were tested in an independent set of globally representative MTB isolates with public sequence and drug resistance data. The validation set showed a higher level of genetic diversity with 44.3% of the 792 isolates belonging to lineage 2, 40.3% belonging to lineage 4 (15% 4.1 sublineages, 8% 4.3 sublineages) and a higher representation of other lineages: 5% L1, 4% L3, 3% L6/BOV/AFR. The proportion of isolates that were MDR in the validation set was 35% (278 isolates). Second line drug resistance phenotypes were available for 25%-57% of the isolates (Table S4) and 29 isolates were XDR. Of the 50 loci identified above, 6 could not be validated as there was no appreciable variation observed in the set of 792 isolates (AF<0.01). Twenty seven other loci were tested but had an AF <0.05 and were not significantly associated, these included the loci *mymA* and *fprA*. Of the remaining 17 loci, 12 were validated to be associated with resistance to one or more drugs. These included *whiB6, ccsA, ubiA*, a metal beta-lactamase *Rv2752c*, and two intergenic regions including *thyX*-*hsdS.1* (Table 1). In the site level analysis the D397G SNV in the gene Rv3805c (*aftB*) was validated as significantly associated with resistance. The strength of association for several of the novel loci was comparable to some canonical genes, but the allele or burden frequency was lower for most of them. For example the effect of *ubiA* mutations on the EMB MIC was measured to be 0.52 logMIC increase, similar in magnitude to the effect of variants in the *Rv1482c-fabG1* intergenic region on INH MIC (0.63 logMIC increase) as was the effect of *whiB6* mutations on SLI MICs (ranging between 0.56–0.60 logMIC increase). The respective allele frequencies were 0.07 for *ubiA*, 0.03–0.04 for *whiB6* and 0.10 for the *Rv1482c-fabG1* intergenic region. The allele frequency in all but two validated loci was <10% (Table 1, Table S2).

All of the validated regions were found to have variants in two or more of the major TB lineages, and all but four of the coding loci harbored nonsense or frameshift variants in one or more isolates (Table S5, Figure S3). The distribution of variants varied by locus; *ubiA, whiB6, Rv2752c and PPE35* all displayed considerable diversity of variants that were closely spaced in one or more segments of the gene, in a pattern similar to that observed in known resistance genes (Figure S3). For the two intergenic hits, variants were most frequent in a distal portion of the region adjacent to the next coding region.

We examined the proportion of variance in the resistance phenotype explained (PVE) by all of the observed genetic variation for each drug (Table 2). The PVE varied by drug, ranging from 0.64 +/- 0.06 for MXF and 0.66 +/- 0.04 for PZA at the lower end to 0.84 +/- 0.02 for RIF and 0.88 +/- 0.02 for AMI at the higher end. We measured the PVE for the known antibiotic resistance genes, and that for novel genes captured in this study. The proportion explained by the known genes was relatively low and at most 0.24 +/- 0.08 (27% of the total PVE) for AMI. The proportion explained by the novel genes was even lower but on par with PVE of known drug resistance loci for PZA and ETA albeit with large error margins (Table 2).

**Table 2:** PVE for each drug attributable to all measurable genetic variation, and those within known drug resistance regions and novel regions associated in this study. Drug abbreviations detailed in Table S7. DR: drug resistance regions as detailed in the methods. Novel: regions specified in Table 1. wo: without.

|  | **All** |  | **wo DR** |  | **wo DR wo Novel** |  | **<u>DR related</u>**<br>**PVE** | **<u>Novel loci</u>**<br>**related PVE** |
| --- | --- | --- | --- | --- | --- | --- | --- | --- |
| **<u>Drug</u>** | **<u>PVE</u>** | **<u>SE</u>** | **<u>PVE</u>** | **<u>SE</u>** | **<u>PVE</u>** | **<u>SE</u>** |  |  |
| **INH** | 0.809 | 0.020 | 0.732 | 0.032 | 0.723 | 0.029 | 0.08 | 0.01 |
| **RIF** | 0.838 | 0.017 | 0.701 | 0.034 | 0.692 | 0.033 | 0.14 | 0.01 |
| **RFB** | 0.833 | 0.023 | 0.722 | 0.041 | 0.693 | 0.042 | 0.11 | 0.03 |
| **EMB** | 0.748 | 0.027 | 0.674 | 0.036 | 0.665 | 0.035 | 0.07 | 0.01 |
| **PZA** | 0.659 | 0.038 | 0.634 | 0.044 | 0.602 | 0.043 | 0.03 | 0.03 |
| **KAN** | 0.833 | 0.022 | 0.671 | 0.040 | 0.658 | 0.041 | 0.16 | 0.01 |
| **AMI** | 0.879 | 0.019 | 0.639 | 0.057 | 0.640 | 0.055 | 0.24 | <0.01 |
| **CAP** | 0.743 | 0.030 | 0.690 | 0.038 | 0.666 | 0.038 | 0.05 | 0.02 |
| **ETA** | 0.701 | 0.034 | 0.689 | 0.038 | 0.675 | 0.038 | 0.01 | 0.01 |
| **STR** | 0.710 | 0.033 | 0.604 | 0.047 | 0.601 | 0.045 | 0.11 | <0.01 |
| **MXF** | 0.643 | 0.058 | 0.494 | 0.083 | 0.456 | 0.081 | 0.15 | 0.04 |

We hypothesized that because antibiotic resistance arises as a result of strong positive selection in MTB that the MIC distributions observed for the same resistance mutation would not vary appreciably across different lineages. We focused on lineage 4 and lineage 2, as they were well represented in our sample. Examining the five mutations: *katG* S315T, *rpoB* S450L, *embB* M306V, *inhA*-15, *pncA* H51R for INH, RIF, EMB, ETA and PZA respectively, we found the MIC distributions not to be appreciably different by the Kolmogrov-smirnov test (P-value >0.5 in all four cases).

Given the number of intergenic regions found to be associated with resistance we tested the hypothesis that intergenic variants have smaller effects on drug MIC compared with gene body mutations for the three genes and the promoter-containing upstream intergenic region that were independently associated with resistance in the GWAS; namely *inhA, pncA* and *embB* and their upstream intergenic regions respectively. We focused on the codon and promoter site with the largest allele frequency in each case. Isolates not infrequently had both a gene body and a promoter mutation: 12% of isolates with *embB* promoter mutations also had an *embB* codon 306V, and 19% of isolates with an *inhA* promoter mutation also had a mutation at *inhA* codon 21. No isolates had both a *pncA* promoter mutation and a *pncA* mutation at codon 51. Figure 2 shows the marginal MIC values for each site pair and drug. Variants in promoter regions consistently showed lower MICs than gene body mutations, although in most cases both medians were above the clinical cutoff (Wilcoxon rank sum test p values 0.03, 0.002, 0.01, 0.009 for the drugs INH, ETA, PZA & EMB respectively). This findings were also supported by the relative magnitude of the GWAS regression coefficients at the locus level for each drug (Table 1) with the notable exception of ETA (Table S2).

**Figure 2:**
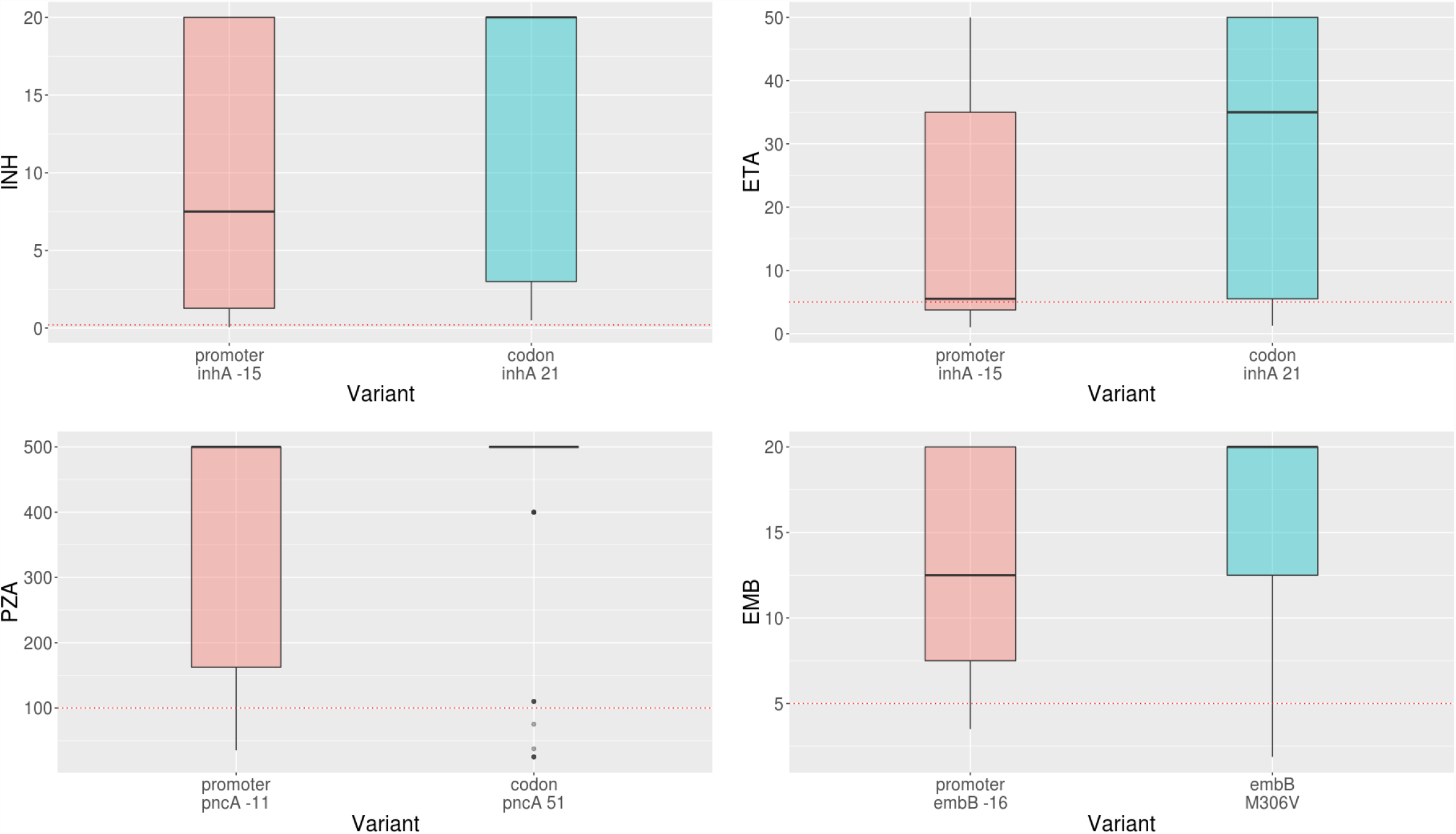
Promoter vs gene body mutations and their effect on MIC (y-axis in ug/mL) for 4 drugs. Wilcoxon rank sum test comparing each pair was significant at p<0.01. For reference critical resistance testing concentration is 0.2ug/mL for INH, 5ug/mL for ETA, 100ug/mL for PZA, 5ug/mL for EMB and indicated by the horizontal dotted line in the box and whiskers plot.

## Discussion

Here, we examine 1452 clinical MTB isolates, enriched for phenotypic resistance, and quantify their antibiotic resistance phenotype using the minimum inhibitory concentration method. Our GWAS results using this quantitative phenotype are notable for the capture of several non-coding genetic regions. In aggregate, more than 20% of the loci associated with antibiotic resistance were intergenic regions. This stands in contrast to the relatively low proportion of the MTB genome annotated to be noncoding, 10.5% by length for the H37Rv reference. Although only a subset of these regions are known promoter regions, their association with higher levels of antibiotic resistance, and the concentration of the variants adjacent to nearby open reading frames raises the possibility that the novel regions may also play a role in gene regulation. Canonically, antibiotic resistance is caused by inactivating protein mutations in drug targets or in pro-drug to drug converting enzymes in MTB. Also, to date, commercial based assays for detecting antibiotic resistance in TB have largely focused on gene based variants, with the notable exception of the *inhA* and *eis* promoters. We find that isolates harboring mutations in promoter regions tend on average to have lower drug MICs than those isolates with a corresponding nonsynonymous gene body variant, and although these tend to exceed the critical cutpoint in both cases, if the MICs are close enough to the cutpoint the isolates may be treatable in some cases with higher doses of drug or a more potent drug from the same class^42,43^. This highlights the importance of understanding the underlying genetic cause of resistance and personalizing therapy based on this, but definitely requires further investigation including potentially clinical trials exploring the efficacy of higher dose antibiotic therapy in patients with such isolates.

We identify and validate 12 genetic regions and one SNV as associated with resistance in *Mycobacterium tuberculosis*. Although these loci have, to date, not been used to predict or diagnose antibiotic resistance in patients with TB^16,17,21^, several have been recently associated with resistance either *in vitro* or in other genome wide association studies performed on binary resistance read outs (Table S6)^11–13^. We summarize these by drug or class, detailing the full results in Table 1 and Tables S2–7.

*Ethambutol*: The gene with the most significant p-value in the primary GWAS was *ubiA*. This locus is validated further by the results of two prior GWAS studies^13,40^, and mutations introduced at *ubiA* codon 237 were shown to increase gene function and elevate decaprenylmonophosphoryl-B-D-ribose or arabinose (DPA) levels^44^. DPA is the donor substrate for arabinosyltransferases that include EmbB, the main target of the drug EMB, and increases in DPA levels likely result in competitive inhibition of the EMB drug effect measurable as an increase in the EMB MIC. The downstream gene *aftB* that encodes an enzyme catalyzing the final step in arabinoglycan arbinan biosynthesis was also found to have a SNV significantly associated with resistance in our study. The association was not with EMB but rather with the drug AMI, as most AMI resistance isolates are also EMB resistant we suspect this mutation to be compensatory to EMB resistance rather than resistance causing, reinforcing *aftB* to be a potentially valuable drug target as has been previously suggested^45,46^.

*Multidrug resistance & pyrazinamide:* The gene Rv2752c encodes a bifunctional beta-lactamase /ribonuclease^47,48^, and was found to be associated with resistance in one prior survey^40^. We found this gene to be associated with resistance to either INH or RIF, with an effect size comparable to that of *inhA* promoter mutations on INH resistance, but with an allele frequency that was half of that of *inhA* promoter mutations (at 0.05). The integral membrane transport protein *KefB* (Rv3236c) is a K^+^/H+ antiporter that releases K^+^ to the phagosomal space and prevents its acidification. We found variants in the encoding gene to associate with resistance most strongly with PZA resistance which is compelling given the known modulating effect of the medium’s pH on PZA’s drug activity^49^.

*Aminoglycosides (AG):* We found several novel associations most strongly with the AG class of anti-tuberculosis drugs. These include the transcriptional regulator whiB6 that is known to activate expression of the DosR regulon, and controls aerobic and anaerobic metabolism and virulence among other pathways^50^. Previous work has implicated another *whiB*-like transcriptional regulator, whiB7, in resistance to AGs^51^ and whiB6 and the upstream intergenic region were previously associated with resistance in a prior GWAS albeit to non-AG agents. The cytochrome-c maturation gene *ccsA* encodes an integral membrane protein that binds heme in the cytoplasm and exports it to the extracellular domain of *ccsB* that in-tail primes it for covalent attachment to apocytochrome *c*. Deficient cytochrome c oxidase activity is tolerated in MTB due to the flexibility of its electron transfer chain^52^, it is plausible that this may incur a fitness advantage by slowing growth under drug pressure.

*Ethionamide:* The gene *mymA*, an alternative monoxygenase to *ethA*^41^ which encodes an enzyme known to activate the prodrug ETA, was associated with an increase in ETA MIC. In vitro, *mymA* deletion mutants were previously found to be resistant to ETA, and double *mymA* and *ethA* knock out mutants had even higher ETA MICs that the individual mutants^41^. We were not able to validate *mymA* in the independent dataset against the binary ETA resistance phenotype, possibly due to limited statistical power as only 116 isolates in the validation dataset where ETA resistant, and *mymA* variants are more rare occurring in <5% of the isolates in the test set. It is also possible that *mymA* mutations increase ETA MIC to a smaller extent in clinical isolates making the GWAS against binary phenotypes less sensitive. The diagnostic utility of *mymA* mutations for improving the prediction of ETA thus requires more study.

*Para-aminosalicylic acid (PAS):* Mutations in the intergenic region upstream of *thyX* (*thyX-hsdS.1*) have been shown to modulate *thyX* expression and have been associated with resistance in two MTB GWA –studies^13,40^. Given that *thyX* is involved in folate metabolism, mutations in these regions may be causative of or compensatory for PAS resistance^13^. It is notable that the association we measured was with respect to other drugs, INH, KAN and AMI, as we did not have a sufficient number of isolates tested for PAS resistance. This likely resulted from drug-drug resistance collinearity and emphasizes the need to carefully interpret novel GWAS results in MDR-bacteria.

The measurement and genome wide association with minimum inhibitory concentrations allowed us to quantify, for the first time, the proportion of the TB resistance phenotype that is explained by bacterial genetic variation. We estimate that 64–88% of the MIC variance to be explained by genetic effects, with standard errors ranging from 2–6%. The remaining proportion may be explained by other factors such as genetic interactions, mutation heterogeneity or environmental or other testing related factors that result in MIC level variability. It is notable that we found the known resistance loci to explain a relatively low amount of the total variation ranging as low as 0.01 for ETA to 0.24 for AMI. The gap between total PVE and that attributable to known drug resistance loci, is not completely explained by the presence of the novel genetic loci as these explained an even lower proportion than known drug resistance loci, likely related to their low mutation frequency. This gap may be better explained by lineage or gene-gene interactions. Although we did assess the interaction between 5 canonical resistance mutations and genetic lineage (lineage 4 vs 2) we could not measure an appreciable interaction for these mutations. It remains possible that such interactions exist for other mutations, especially those with smaller effects as these have been noted previously in allelic exchange experiments^53^.

In this study, we demonstrate the utility of genome wide association for examining bacterial phenotypes relevant to infectious disease. Our study was not without limitations. Given the recognized step wise acquisition of resistance in MTB^54^, it is very challenging to determine accurately which drug resistance is in fact associated with a particular gene or genetic region. For example resistance to any of the second line agents, fluoroquinolones like MXF, SLI’s like AMI, or to first line agents like PZA and EMB, nearly always co-exists with resistance to INH and RIF, and it is thus not possible to perform association conditioning on the absence of resistance to those agents. Further, the performance of linear mixed models for performing GWAS in bacteria has not been systematically studied, although applied recently to MTB and other bacteria with demonstrated success^13,55–57^. We acknowledge that we cannot be certain that these models adequately control for population structure in clonal bacteria and because of this we performed validation in an independent dataset with a different lineage distribution. We also provide the lineage breakdown of variants in our hit loci that in each case demonstrated evidence for convergent evolution^11^. We also demonstrate the power of using a binary gene-burden score for bacterial GWAS, as this decreased the number of necessary tests relative to GWAS of individual sites and allowed the incorporation of rare genetic variants that appear to be important for drug resistance in MTB^58^. This approach is however, reliant on the accuracy of the available genomic annotation for MTB, and is most sensitive for capturing genes under diversifying selection, i.e. where multiple different genetic mutations may contribute to a functional genetic change, and entirely ignores synonymous variation as potentially contributing to the phenotype. More refined measures of gene burden in bacteria, for example measures that incorporate protein structural data, are worth investigating systematically in the future.

In summary, with the increasing availability of genomic data, powered by the formation of TB genomic data consortia^38^, our ability to identify more rare variants with smaller effects on resistance will increase. Our improved understanding of the genetic mechanisms of resistance in MTB can perhaps lead to more targeted drug development efforts, but more imminently will allow for improved diagnosis and surveillance given the increased uptake of genomic technologies in public health laboratories in high income countries^59^. Improvement in portable sequencing technology^60^ and decreased cost of sequencing is promising to facilitate adoption in settings with lower resources were TB is most prevalent. However even if sequencing technology is available, our results suggest that genomic data interpretation will likely necessitate the use of statistical models or machine learning^16,58,61^ given the number of genetic loci associated with resistance and the likely contribution of gene-gene interactions, especially if a quantitative prediction of the drug MIC is desirable^15^. The portability of the potential benefits of these advances to areas of the world where TB is most prevalent will require continued efforts in open sharing of data and analysis tools^62^.

## Online Methods

### Sample Collection

MTB sputum based culture isolates were selected from (1) a Peruvian patient archive of culture isolates enriched for resistance based on prior targeted resistance gene sequencing and binary DST phenotype^16^ (n=496), or (2) sampled from a longitudinal cohort of patients with Tuberculosis from Lima Peru^18^ enriched for multidrug resistance based on prior binary DST (n=568). These 1064 isolates had phenotypic resistance testing by MIC for 12 drugs repeated (see below) at the National Jewish Hospital (NJH) Denver, CO, and underwent whole genome sequencing. Data from these isolates were pooled with data from two additional samples: a convenience sample from three national or supranational reference laboratories selected based on the availability of MIC data: the Institute for Tropical Medicine -Antwerp, Belgium, the Massachusetts State TB Reference Laboratory -Boston, MA, and the National Institute for Public Health and the Environment -Bilthoven, Netherlands (n=411) and a sample of 83 pan-susceptible isolates from the Peruvian TB cohort^18^ added to increase the representation of sensitive isolates.

### Culture and Drug resistance/MIC testing

Lowenstein-Jensen (LJ) culture was performed from sputum specimens using standard NALC-NaOH decontamination. Prior to DNA extraction and sequencing most cultures had been cryopreserved as follows: Inside a biosafety container, all colonies of each culture were extracted from the LJ slants and dissolved in 7H9 broth with 20% glycerol to reach a bacterial suspension similar or higher than McFarland 5. Then the bacterial suspension was aliquoted in volumes of 0.3 to 0.5 mL and stored overnight at 4° C to ensure the glycerol uptake of the cells. Then, all tubes were placed into the –80° C freezer for long term storage.

All isolates, except the 83 pan-susceptible isolates described above, underwent minimum inhibitory concentration testing. Testing for the 1064 isolates at NJH was performed for 12 anti-TB drugs on 7H10 media using agar proportion in a staged fashion. Isolates were first tested at three low concentrations that include the WHO recommended critical concentration. If the isolate was resistant at the critical concentration then testing at six higher concentrations was additionally performed. The testing concentrations deviated from the traditional doubling to better detect intermediate level MICs that are close to the clinical critical concentration and within theoretically achievable levels in patient sera based on available pharmacodynamics data^19^. The concentrations are detailed in Table S7. Culture, MIC and DST testing at the other labs is outlined in Table S8. Testing methods and concentrations are also listed for each isolate in Table S1.

### DNA extraction and Whole genome sequencing

DNA from sputum samples of TB patients was extracted from cryopreserved cultures. Each isolate was thawed and subcultured on LJ and a big loop of colonies were lysed with lysozyme and proteinase K to obtain DNA using CTAB / Chloroform extraction and ethanol precipitation. DNA was sheared into ∼250bp fragments using a Covaris sonicator (Covaris,Inc.), and prepared using the TruSeq Whole-Genome Sequencing DNA sample preparation kit (Illumina, Inc.). Samples were sequenced on an Illumina HiSeq 2500 sequencer. Paired-end reads of length 125 bp were collected. Base-calling was performed using HCS 2.2.58 and RTA 1.18.64 software (Illumina, Inc.)

### Definition of known drug resistance loci

We define the MTB known resistance loci as the following genes *katG, inhA* & its promoter, ahpC promoter, *kasA, rpoB, embA, embB, embC* & *embA-embC* intergenic region, *ethA, gyrA, gyrB, rrs, rpsL, gid, pncA* & its promoter, *tlyA, thyA, rpsA, eis* promoter and the compensatory genes *rpoC, rpoA* based on prior published work^6,16,20–25^.

### Variant calling and phylogeny construction

We aligned the Illumina reads to the reference MTB isolate H37Rv NC_000962.3 using Stampy 1.0.23^26^ and variants were called by Platypus 0.5.2^27^ using default parameters. Genome coverage was assessed using SAMtools 0.1.18^28^ and FastQC^29^ and read mapping taxonomy was assessed using Kraken^30^. Strains that failed sequencing at a coverage of less than 95% at ≥10x of the known drug resistance regions, or that had a mapping percentage of less than 90% to *M. tuberculosis* complex were excluded. Genomic regions not covered at ≥10x in at least 95% of the remaining isolates were filtered out from the analysis, i.e. no attempt at association with variants in those regions was made. In the remaining regions, variants were further filtered if they had a quality of <15, purity of <0.4 or did not meet the PASS filter designation by Platypus. We also excluded any indels >3bp in size or large sequence polymorphisms. Further quality control was performed after genome wide association when associated PE/PPE gene and indels were visualized and manually inspected using IGV v2.4.9^31^. TB genetic lineage was called using the Coll *et al*.^32^ SNP barcode and confirmed by constructing a Neighbor joining phylogeny using MEGA-5^33^ including lineage representative MTB isolates from Sekizuka *et al*.^34^.

### Genome wide association & Validation

*Phenotype:* The MIC data was recorded as an interval indicating the last highest concentration tested where growth was seen and the MIC itself. Because critical concentrations on LJ media (for isolates tested at ITM) are in general higher than those on 7H10, the MIC intervals were normalized to allow for comparability by dividing by the critical concentration for each drug as defined by the WHO^35^. The interval midpoints were computed and converted to ranks as has been previously suggested for genotypic association with MIC data^36^; ties were assigned an average rank. A sensitivity analysis was performed to confirm that the results are not sensitive to the rank transformation of the phenotype, by comparing the region hits obtained in a parallel GWAS analysis using the natural log transformed phenotype instead of the rank transform.

*Genotype & GWAS:* Association analysis was performed at the gene/non-coding region level using a binary gene burden score that was set at ‘one’ if any non-synonymous single nucleotide substitution (SNV) or indel (insertion or deletion) was observed in a gene, or any SNV or indel was observed in a non-coding region, and ‘zero’ otherwise. We excluded known lineage markers in drug resistance genes from the burden score calculation^16^. Association was also performed at the site level in a secondary analysis excluding synonymous variants. Any gene/region or SNV with a minor allele frequency (MAF) of <0.01 was not tested. We controlled for population structure by computing a genetic relatedness matrix (GRM), including all synonymous and non-synonymous SNVs & indels but excluding variants in known drug resistance loci and variants occurring at a MAF of <0.01 using the software package GEMMA^37^. Genome wide association was performed using a linear mixed model with the phenotype as the rank-transformed MICs also using GEMMA. Regions with a false discovery rate <0.05 were selected for validation. We verified control for population structure with QQplots using the qqman package in R v3.2.3. As the regression was performed on rank transformed MIC values, we scaled the resulting effect size back to the MIC scale by first performing a linear regression between the natural log MIC values and their rank transform and then using the resulting slopes as a scaling factor. LogMIC change in units of log(mg/L) are reported throughout.

*Validation:* We validated the genomic regions identified above in an independent public dataset with binary phenotype data. The validation dataset consisted of a convenience sample of 792 MTB isolates obtained by pooling data from the ReSeqTB knowledge base (https://platform.reseqtb.org/)^38^ with additional MTB whole genome sequences and phenotype data curated manually from the following references^39,40^ (Table S4). We did not select isolates for the validation set based on lineage or drug resistance profiles. Association analysis was performed only using a similar linear mixed model approach as was outlined above using a GRM for population structure correction. A locus was considered validated if it had a Wald p-value of <0.005.

*Proportion of variance explained (PVE):* We computed the PVE as the proportion of total phenotypic variance explained by the genetic relatedness between the isolates, using the restricted maximum likelihood approach as implemented in GEMMA, as a measure of heritability. We computed the PVE attributable to known drug resistance regions by recomputing the GRM after removing all variation (synonymous, nonsynonymous and indels) in the known resistance loci. Similarly we computed the PVE attributable to all other loci validated to be significantly associated with resistance in this study, as PVE attributable to ‘novel’ loci. Given the phenotypes were coded as ranks of the MIC distribution, we performed a sensitivity analysis to confirm that rank transformation did not affect our PVE measurements. In this sensitivity analysis we dichotomized the MICs using the WHO established critical concentration as the threshold, and recomputed the PVE on the liability scale. The PVEs changed by <10% for all drugs in the sensitivity analysis.

## Data

All data used in this study is available in the supplementary material or deposited on NCBI with accession numbers detailed in Table S1.

## Acknowledgement

MF, and MM conceived this study, MF conducted the analysis and wrote the first version of the paper with key input from all authors. LF provided analysis support, and SS and MM provided analysis oversight. BdJ, LR and CJM curated, phenotyped and sequenced the TDR isolates. AS and DK curated tested the isolates from MSLI. DvS curated and phenotyped the isolates from RIVM. JS and MS sequenced the isolates from MSLI and RIVM. RC cultured and maintained the archive of isolates from SES. JS and TI performed the sequencing and quality control on all SES isolates. We would like to acknowledge the TB patients and their providers who provided the samples for this this study and without which it would not have been possible. We acknowledge the ReseqTB team (Drs. Marco Schito and Matthew Esmundo for providing us with data that allowed the validation of our GWAS hits).

